# Doublet decoding of tRNA^Ser3^ demonstrates plasticity of ribosomal decoding center

**DOI:** 10.1101/2024.12.14.628468

**Authors:** Shruthi Krishnaswamy, Shirin Akbar, Daniel S. D. Larsson, Yang Chen, Maria Selmer

## Abstract

Frameshifts can be caused by specific combinations of tRNA and mRNA. The wildtype AGC-decoding *E. coli* tRNA^Ser3^__GCU__ was in the 1980s shown to induce -1 ribosomal frameshifting on GCA alanine codons, and proposed to read a two-base codon instead of a canonical triplet. It has remained unclear whether this type of non-cognate decoding can be accommodated by the ribosome. We here performed single-particle cryo-EM reconstructions on *E. coli* 70S ribosomes with the frameshift-inducing tRNA^Ser3^ bound to the non-cognate GCA codon or the cognate AGC codon in the ribosomal A site. The structures demonstrate that doublet-decoding is made possible when A1493, the conserved monitoring base in 16S rRNA, mimics a first codon base, forming a Hoogsteen base pair with U36 from the anticodon and stacking with the mRNA. This interaction pushes the first two bases of the A-site codon in position for base pairing with C35 and G34 of the anticodon.

## Main text

The ribosome has evolved to assure high fidelity translation of the genetic code and maintenance of the three-base reading frame (reviewed in^1,2^). The decoding center of the ribosome monitors the geometry of the base pairs between the aminoacyl tRNA and the A-site codon. This is coupled to the closure of the 30S ribosomal subunit that triggers GTP hydrolysis of EF-Tu, leading to its dissociation, followed by accommodation of the aminoacyl tRNA and peptidyl transfer. While the A-site is empty, the ribosomal interactions with P-site tRNA maintain the reading frame^3^. Several mechanisms have been described for regulatory and erroneous frameshifts. Most of these involve slippage of the P-site tRNA in response to pausing or strong mRNA structure, but some occur at the ribosomal A-site. Such frameshifts can *e*.*g*. be induced by mutated or under-modified tRNAs (reviewed in^4,5^).

In 1979, Atkins *et al*. observed, in a cell-free translation system from *E. coli* translating the MS2 coat protein gene, that a non-cognate tRNA^Ser3^__GCU__ could promote -1 frameshifting on GCA Ala codons^6^. Similarly, tRNA^Thr3^__GGU__ could induce frameshift on CCG/A codons ^6^. Such infrequent shifts to the -1 reading frame were proposed to have regulatory roles in phages MS2 and ϕX174 ^7,8^. Further experiments showed that the frameshifting propensity of tRNA^Ser3^ only resided in the anticodon loop, and that this property could be transferred to the body of tRNA^Phe^. This led to proposal of the doublet-decoding hypothesis^9^, suggesting that the anticodon loop of tRNA^Ser3^ would adopt a conformation where a two-base anticodon (G_34_C_35_) could read only the first two bases of the GCA Ala codon (Figure 1a), establishing a new reading frame already in the A site. A later suggestion was that altered stacking in tRNA^Ser3^_GCU_ would allow presentation of a shifted anticodon, with U_33_ forming an additional base pair with the mRNA, in which case the -1 frameshift would instead happen in the P site, when the tRNA would regain its normal conformation^10^. We here set out to test the doublet-decoding and shifted-anticodon hypotheses and investigate if and how the *E. coli* ribosome could accept this variant of non-cognate tRNA.

**Figure 1.**
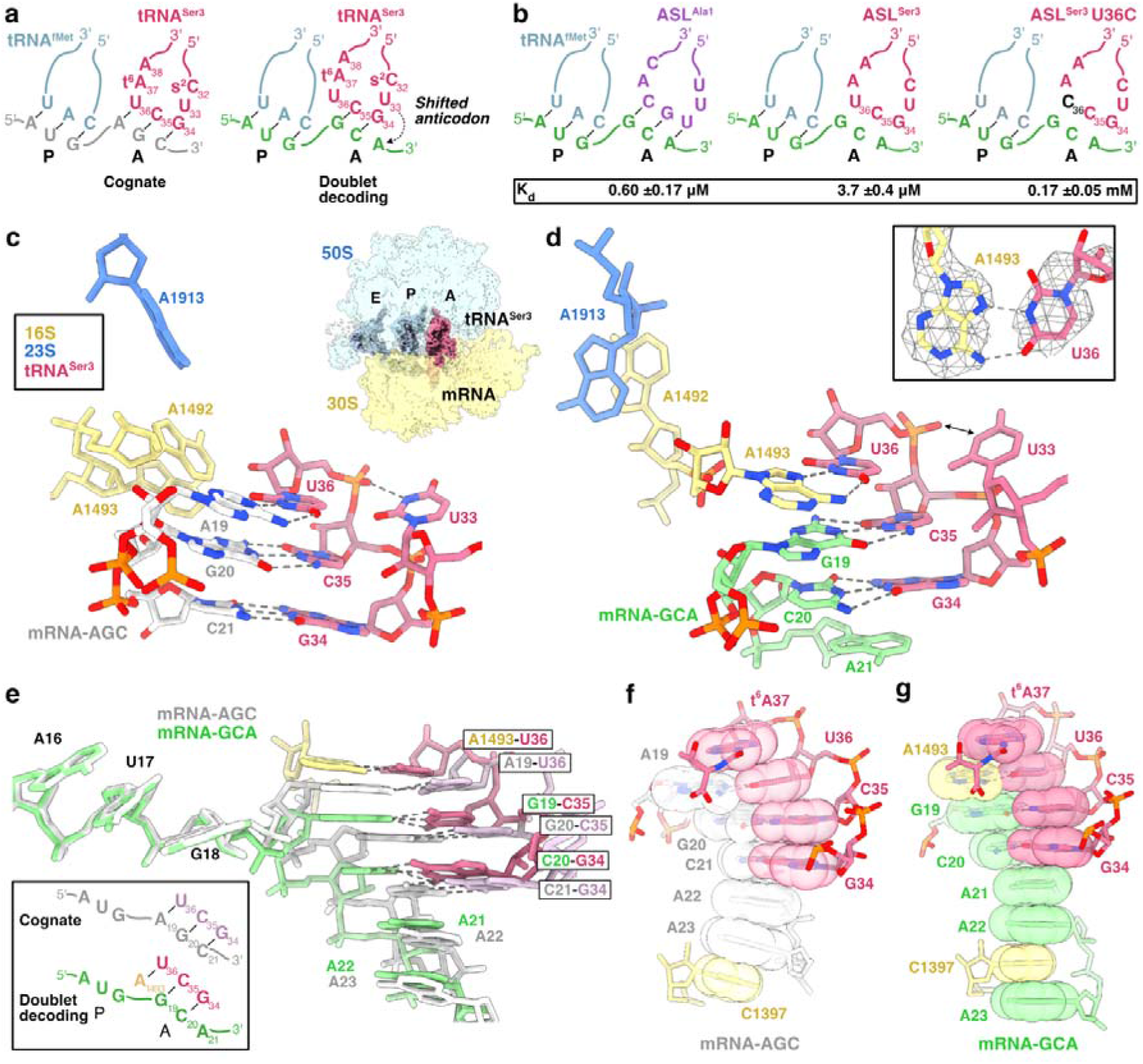
Doublet decoding in the *E. coli* ribosome. **a.** Schematic representation of cognate and doublet-decoding ribosomal complexes with tRNA^Ser3^ in the A-site. **b**. Schematic representation of complexes used in filter-binding experiments, with results in the box. **c**. Structure of the decoding center with cognate tRNA^Ser3^ and mRNA-AGC. Cryo-EM reconstruction of the ribosomal complex is shown as inset. **d**. Structure of the decoding center with doublet-decoding tRNA^Ser3^ and mRNA-GCA. Inset shows the A1493-U36 Hoogsteen base pair in the cryo-EM map (mesh). **e**. Superposition and schematic of cognate mRNA-GAC and doublet decoding mRNA-GCA from the two structures showing displacement of G19 by A1493 in doublet decoding. **f-g**. Codon-anticodon stacks in the cognate and doublet-decoding structures.

To test whether tRNA^Ser3^ would bind to the GCA codon in a minimal system, we used nitrocellulose filter binding to measure the equilibrium binding affinity of anticodon stem-loops (ASLs) corresponding to tRNA^Ser3^ and the cognate tRNA^Ala1^ to the ribosomal A-site of 70S complexes programmed with a short synthetic mRNA-GCA and tRNA^fMet^ (Figure 1b). The frameshift-inducing ASL^Ser3^_GCU_ showed 6-fold lower affinity than the cognate ASL^Ala1^_UGC_ (Figures 1b, S1). For reference, the K_d_ of ASL^Phe^_GAA_ to its two cognate codons UUC and UUU is 7.6 μM and 0.86 μM, with a GC Watson-Crick or GU wobble base pair in the third position^11^. Thus, the non-canonical binding of ASL^Ser3^ to the GCA codon shows affinity similar to a standard codon-anticodon interaction. In line with earlier studies showing that U36 is critical for -1 frameshifting of tRNA^Ser3 9^, ASL^Ser3^ U36C showed a further >40-fold reduced affinity to the GCA codon, equivalent to a non-cognate interaction (Figures 1b, S1). This confirms that the mRNA does not make a -1 shift upon ASL binding to form an A-site GGC codon, allowing C36 to form a base pair.

To capture the structure of the doublet-decoding state, we prepared complexes of *E. coli* 70S ribosomes with the same synthetic mRNA-GCA, tRNA^fMet^ and tRNA^Ser3^ for single-particle cryogenic electron microscopy (cryo-EM) imaging. A cognate control sample was assembled with mRNA-AGC (Figure 1a). The reconstructions yielded maps with global resolution of 2.61 Å for the doublet-decoding complex and 2.49 Å for the cognate complex (Figures S2–4, Table S3).

In the cognate complex, three Watson-Crick base pairs are formed between anticodon bases G34, C35 and U36 and the cognate AGC codon (Figure 1c, S5a), and the monitoring bases A1492, A1493 and G530 of 16S rRNA make the expected interactions^12^ with the base pairs (Figure S5b).

In the doublet-decoding complex, anticodon bases G34 and C35 make Watson-Crick interactions with the first two bases of the GCA codon, while U36 forms an unpredicted Hoogsteen base pair with A1493 of 16S rRNA (Figure 1d, S5c). This explains the critical role of U36, as cytosine cannot form a Hoogsteen base pair with adenosine^13^. The base pairing of A1493 prevents A1492 from reaching and interacting with the codon-anticodon pair, and it instead stacks with A1913 of 23S rRNA (Figure 1d, S5d).

Superposition of the two complexes based on the 3’ part of 16S rRNA (Figure 1e) shows that doublet decoding is enabled when A1493 replaces the first codon base, pushing G19, the first A-site base, into position to base pair with C35 of the anticodon. This interaction induces the - 1 frameshift that presumably will be finalized by two-base translocation to the P site. The two tRNA^Ser3^ structures show very similar three base-pair stacks further stabilized by t^6^A37 on top and the following mRNA bases and C1397 of 16S rRNA below (Figure 1f-g), disproving the shifted-anticodon hypothesis. Similar stacking of A1493 or its equivalent with the first position of the A-site codon is observed in several ribosome structures in classical state with a vacant A-site (e.g. ^14,15^ from bacteria and eukaryotes, Table S4), suggesting that doublet decoding occurs by conformational selection followed by local adjustment of the mRNA. The “clamping” of the downstream mRNA stack by A1493 and C1397 or its equivalents has been suggested to contribute to reading frame maintenance and prime the mRNA for tRNA selection^14,15^, but in this special case appears to prime the ribosome for doublet decoding. Detailed comparison shows that during doublet decoding, the anticodon loop is wider around the universally conserved U33, weakening its interaction with phosphate 36 (Figure S6). U33 is critical for cognate decoding^16^, but doublet decoding was shown to tolerate an U33A substitution^9^, which is explained by the lack of canonical U33 interactions in the wider anticodon loop.

The global conformation of the 30S subunit in the cognate complex is predominantly closed (Figure S7a), mediated by G530 interactions with tRNA, mRNA and the monitoring bases (Figure S7b), while the doublet-decoding complex is more open, with G530 unengaged (Figure S7c), as previously observed in near-cognate complexes^17^. Thus, to maintain the essential fidelity of translation, the ribosome will only rarely accept this type of non-cognate, frameshift-inducing tRNA.

The doublet decoding model of -1 frameshift was proposed for tRNA^Ser3^_GCU_ on GCA Ala codons and for tRNA^Thr3^_GGU_ on CCG/A codons ^6^. Our structures reveal that U_36_ is critical for formation of a Hoogsteen base pair with A1493 (Figure 1d), and the two tRNA-mRNA base pairs likely have to be G-C for sufficient affinity. These two base pairs are stacked between A1493 and the third mRNA base A_21_ (Figure 1d, 1g), in both cases involving A and G, which form the strongest stacking interactions^18,19^.

In conclusion, this study confirms the doublet-decoding hypothesis. In line with observations for other types of frameshifting, the frameshift inducer tRNA^Ser3^ exploits universally conserved features of the ribosome^5^, in this case the plasticity of the decoding center, to perturb the reading frame. Future studies will elucidate the presumably low frequency of this phenomenon *in vivo* as well as the sequence and organism limitations for doublet decoding.

## Acknowledgements

Cryo-EM grid preparation and data collection was done at the Cryo-EM Uppsala facility, funded by the Department of Cell and Molecular Biology, the Disciplinary Domains of Science and Technology and of Medicine and Pharmacy at Uppsala University. We acknowledge Athina Eleftheraki, Nour Aldin Kahlous, Tana Tandaric and Liuqun Zhao for grid freezing and feasibility test. This research was funded by grants from the Swedish Research Council (2016-06264 and 2022-04511) and from the Swedish Foundation for Strategic Research (F06-0010) to M.S.

