## Supplementary material for "Doublet decoding of tRNA^Ser3^ demonstrates plasticity of ribosomal decoding center": Table S1-S4 Figure S1-S7

**Content:**

Table S1-S4
Figure S1-S7
Supplementary references

**Table S1.** RNA oligos. Anticodons and A-site codons in bold.

| | |
|---|---|
| ASL$^{Ala1}$ | CCUGCUU**UGC**ACGCAGG |
| ASL$^{Ser3}$ | CUCCCCU**GCU**AAGGGAG |
| ASL$^{Ser3}$ U36C | CUCCCCU**GCC**AAGGGAG |
| mRNA-AGC | GGCAAGGAGGUAAAAAUG**AGC**AAA |
| mRNA-GCA | GGCAAGGAGGUAAAAAUG**GCA**AAA |

**Table S2. DNA oligos.**

| | |
|---|---|
| ser3_exp_f1 (EcoRI) | GGAATTCTGTGGTGAGGTGGCCGAGAGGCTGAAGGCGCTCCCCT GCTAAGGGAGTATGCGGTCAAAAGCT |
| ser3_exp_b1 (PstI) | AACTGCAGGCAAATGGCGGTGAGGCGGGGATTCGAACCCCGGAT GCAGCTTTTGACCGCATACTCCC |

**Table S3. Cryo-EM data collection, refinement and validation statistics**

| | #1 Cognate (EMDB-51758) (PDB 9H0L) | #2 Doublet decoding (EMDB-51679) (PDB 9GXX) |
|---|---|---|
| **Data collection and processing** | | |
| Magnification | 190,000 | 190,000 |
| Voltage (kV) | 200 | 200 |
| Electron exposure (e–/Å$^2$) | 28.5 | 28.3 |
| Defocus range (μm) | -0.7 to -1.3 | -0.7 to -1.3 |
| Pixel size (Å) | 0.7463 | 0.7463 |
| Symmetry imposed | C1 | C1 |
| Initial particle images (no.) | 751,778 | 1,051,022 |
| Final particle images (no.) | 167,987 | 196,921 |
| Map resolution (Å) | 2.49 | 2.61 |
| FSC threshold | 0.143 | 0.143 |
| Map resolution range (Å) | 1.86–43.6 | 1.91–41.8 |
| **Refinement** | | |
| Initial models used (PDB codes) | 7K00, 8CGK, 8CGJ, 8CF1 | 7K00, 8CGK, 8CGJ, 8CF1 |
| Model resolution (Å) | 2.65 | 2.78 |
| FSC threshold | 0.5 | 0.5 |
| Map sharpening *B* factor (Å$^2$) | -64.9 | -67.2 |
| Model composition | | |
| Non-hydrogen atoms | 144,192 | 144,033 |
| Protein residues | 5,587 | 5,587 |
| RNA residues | 4,657 | 4,648 |
| Waters | 0 | 0 |
| $Mg^{2+}$ | 314 | 288 |
| *B* factors (Å$^2$) | 114 | 114 |
| Protein | 122 | 121 |
| RNA | 111 | 111 |
| R.m.s. deviations | | |
| Bond lengths (Å) | 0.009 | 0.009 |
| Bond angles (°) | 1.70 | 1.52 |
| Validation | | |
| MolProbity score | 1.18 | 1.21 |
| Clashscore | 2.35 | 2.91 |
| Poor rotamers (%) | 0.87 | 0.72 |
| Ramachandran plot | | |
| Favored (%) | 97.06 | 97.32 |
| Allowed (%) | 2.90 | 2.66 |
| Disallowed (%) | 0.04 | 0.02 |
| Rama-Z score | -1.13 | -1.15 |

**Table S4. Identified structures where A1493 or equivalent is stacked with the first base of the A-site codon**

| PDB code | Comment | Stack (chain id:base) |
|---|---|---|
| 8P2F[1] | *Staphylococcus aureus* 70S ribosome with elongation factor G locked with fusidic acid cyclopentane in post-translocational state | HA:A1504<br>IA:G19 |
| 4V9A[2] | Crystal Structure of the 70S ribosome with tetracycline. (*Thermus thermophilus*) | A:A1466<br>W:U4 |
| 7N31[3] | Elongating 70S ribosome complex in a post-translocation (POST) conformation (*Escherichia coli*) | A:A1493<br>V:G46 |
| 6GZ3[4] | tRNA translocation by the eukaryotic 80S ribosome and the impact of GTP hydrolysis, Translocation-intermediate-POST-1 (TI-POST-1) (*Oryctolagus cuniculus*, rabbit) | E:A1664<br>C:A8 |
| 7ACR[5] | Structure of post-translocated trans-translation complex on *E. coli* stalled ribosome (i.e. no P-site tRNA). | B:A1493<br>D:G90 |
| 7NWT[6] | Initiated 70S ribosome in complex with 2A protein from encephalomyocarditis virus (EMCV) | 2:A1493<br>XX:G43 |
| 7QP7[7] | Structure of the human 48S initiation complex in closed state (h48S AUG closed) | A:A1825<br>7:G136 |

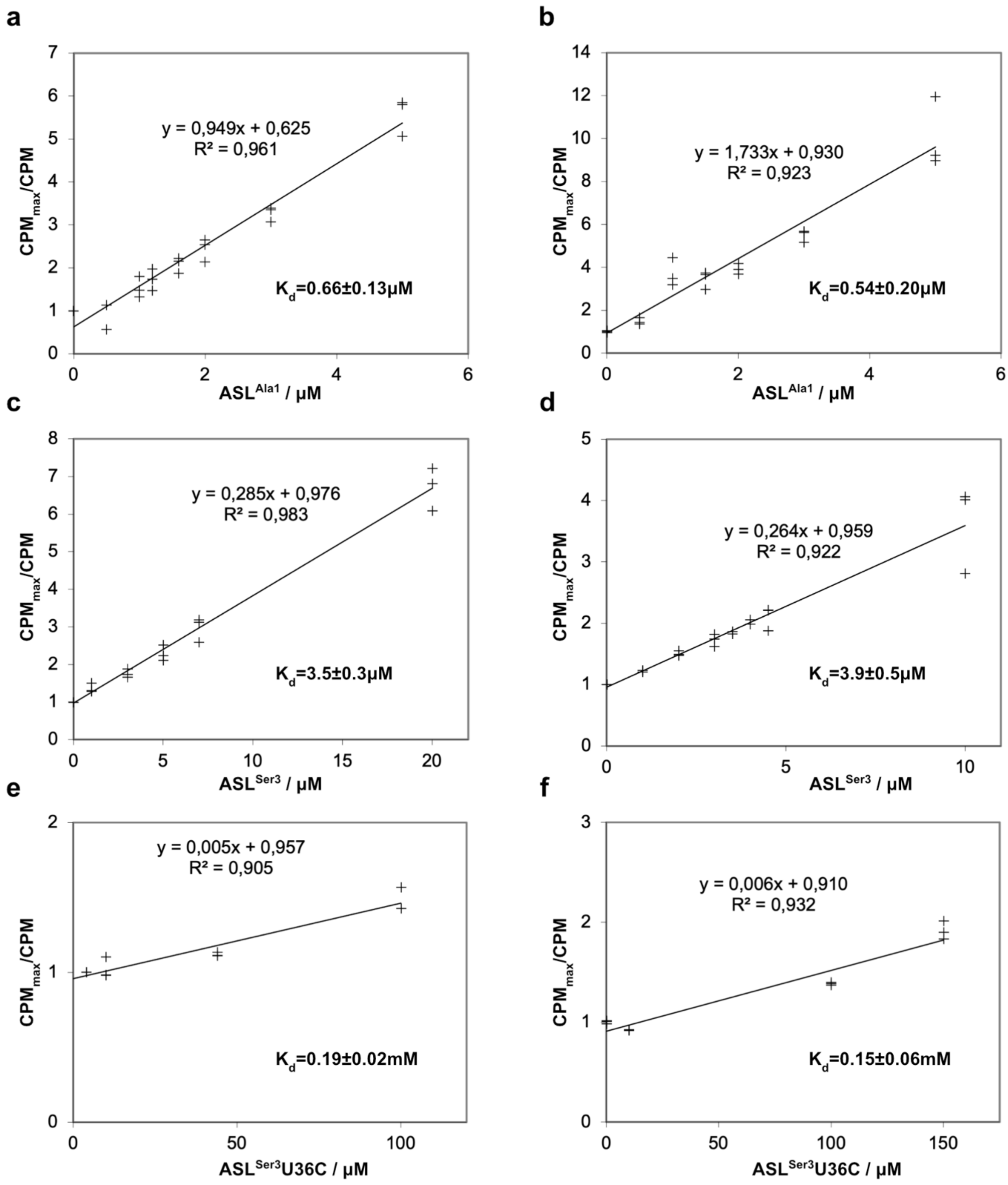

**Figure S1.** Nitrocellulose filter binding curves of $ASL^{Ala1}$ (**a, b**), $ASL^{Ser3}$ (**c, d**) and $ASL^{Ser3}$U36C (**e, f**) to the A-site GCA codon of the *E. coli* 70S ribosome. The X axis shows the concentration of cold competitor ASL and the Y axis the ratio of the background-subtracted CPM values at zero concentration (no competitor) and at the indicated concentration. From this plot, $K_d$ is derived as the ratio between the intercept and the slope of the fitted line. The sum of percentage standard deviation of the intercept and the slope is summed and applied to the $K_d$.

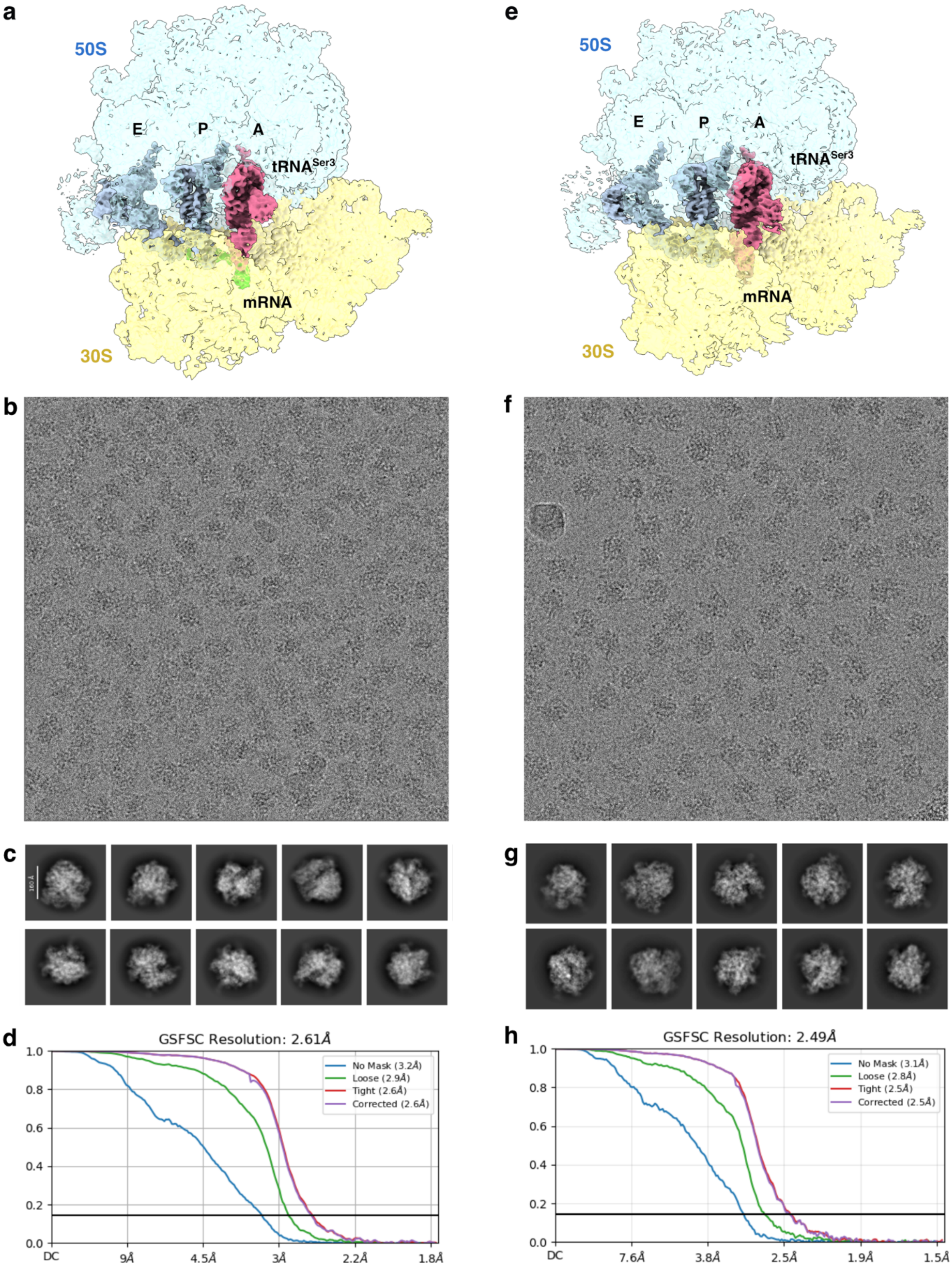

**Figure S2**. Cryo-EM reconstruction, representative micrograph, 2D class averages and FSC plot for cryo-EM reconstructions of doublet decoding (**a-d**) and cognate (**e-g**) complex.

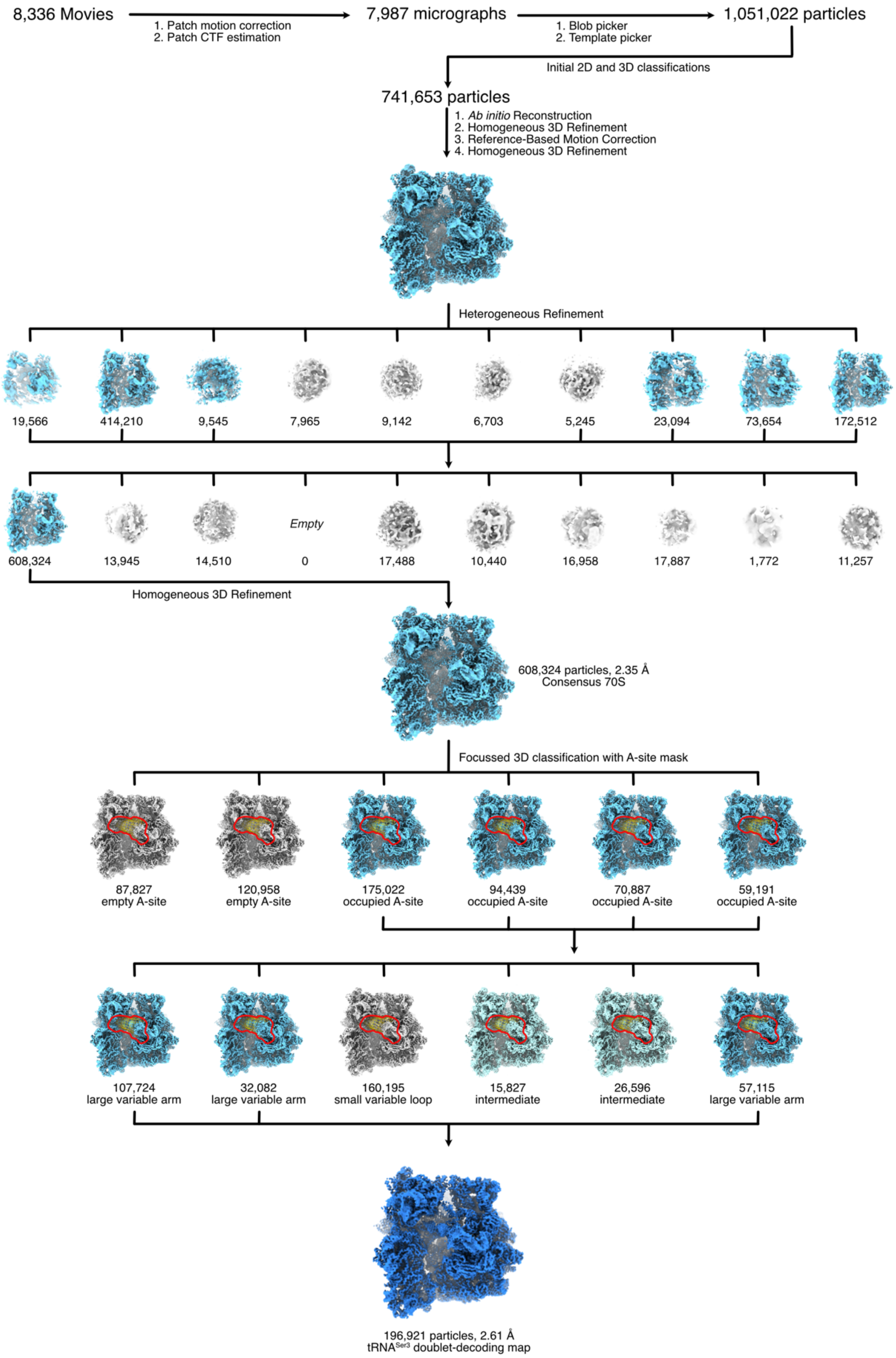

**Figure S3.** Cryo-EM data processing workflow for the doublet decoding structure. Focused mask region for the A-site classification is shown in red outline.

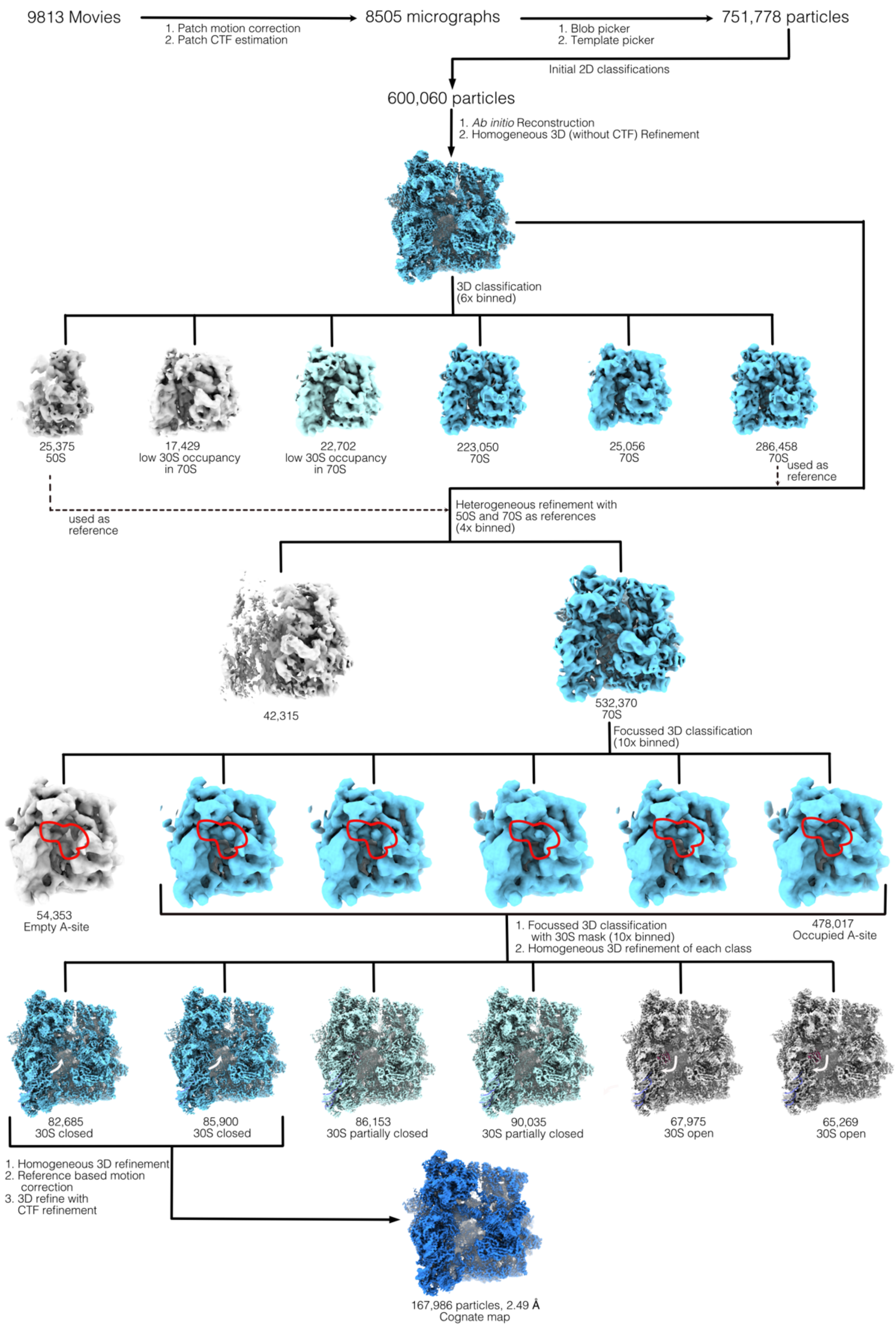

**Figure S4.** Cryo-EM data processing workflow for the cognate structure. Focused mask region for the A-site classification is shown in red outline. 30S conformation shown with inward or outward facing white arrows for closed or open conformation respectively.

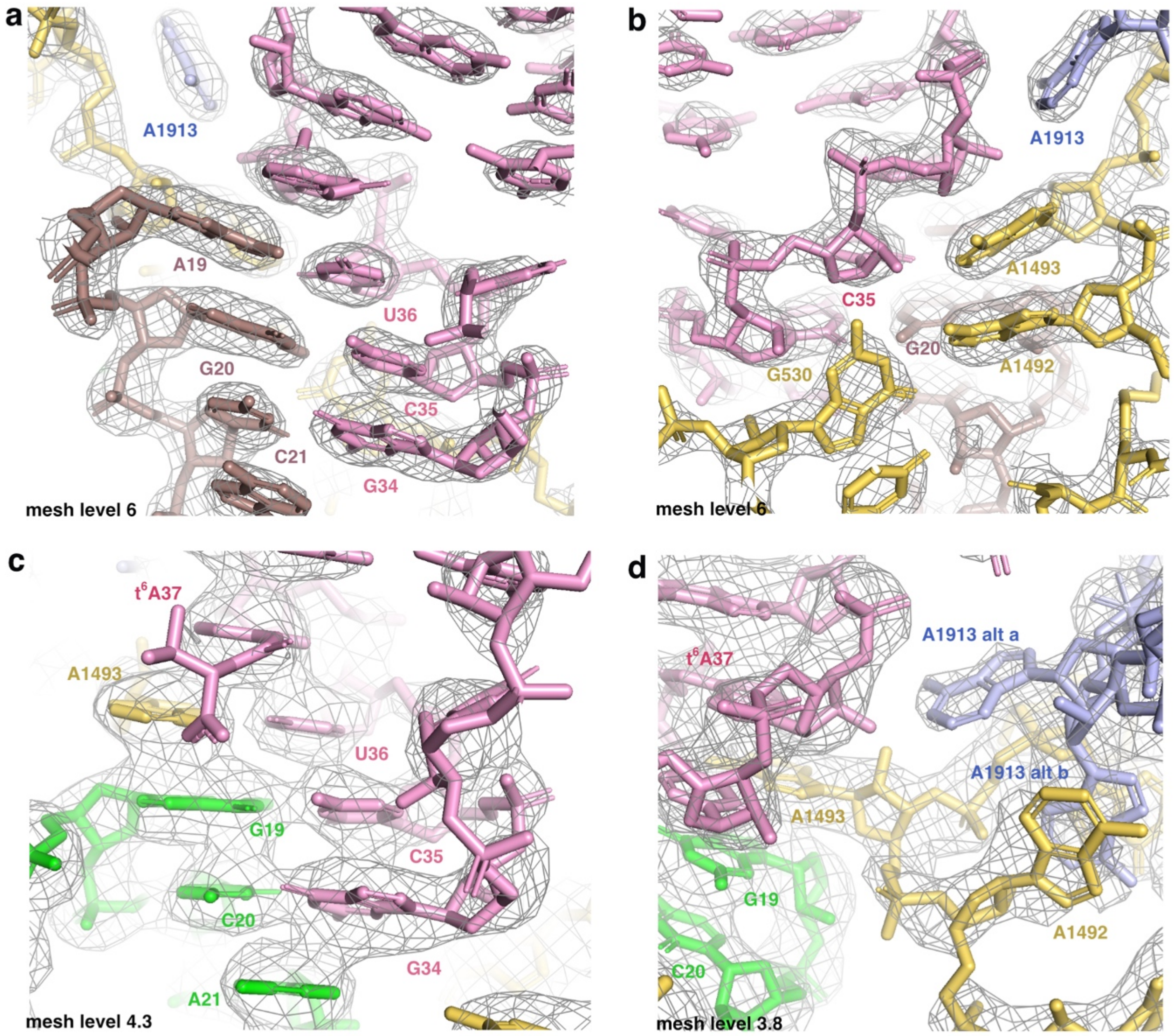

**Figure S5.** Cryo-EM map density shown in mesh with fitted coordinates for the cognate (**a, b**) and doublet-decoding (**c, d**) structures. 30S is shown in yellow, 50S in light blue, tRNA in pink. mRNA-AGC is shown in brown (cognate) and mRNA-GCA in green (doublet-decoding).

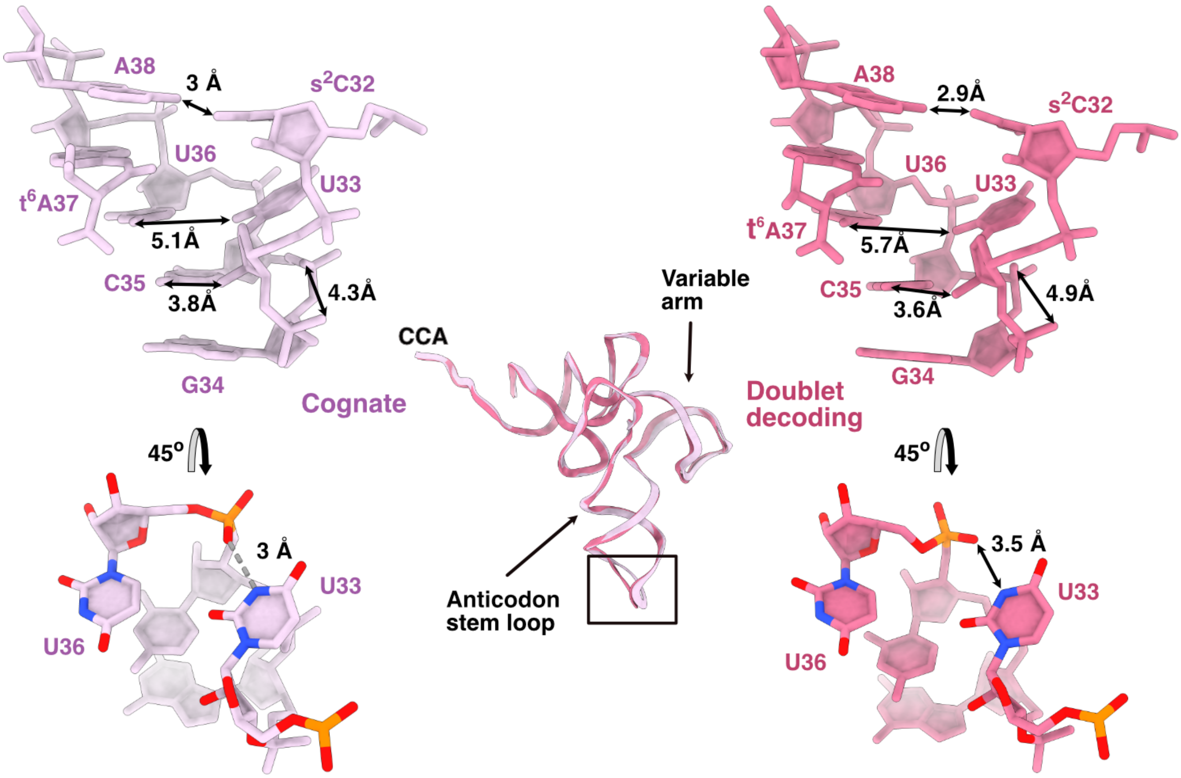

**Figure S6.** Comparison of the anticodon loop of $tRNA^{Ser3}$ at the A-site from the cognate and doublet-decoding complex. Change in distances are shown with double-headed arrows. Lower panel shows increase in distance in the doublet-decoding complex resulting in breaking of hydrogen bond between the U33 base and the U36 backbone phosphate.

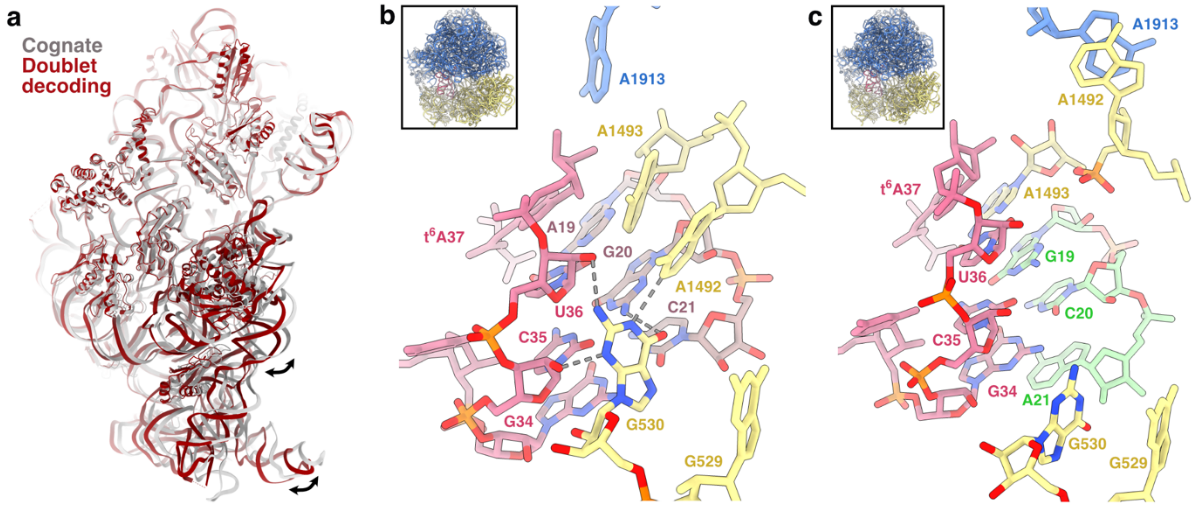

**Figure S7. a**) Superposition of 30S from cognate (gray) and doublet decoding (maroon) structure aligned on the 30S head domain showing the closed and open 30S conformations. Black arrows indicate the direction of movement. **b**) G530 of 16S rRNA in the cognate structure with closed 30S conformation, showing hydrogen bonds to $tRNA^{Ser3}$ (pink), mRNA-AGC (brown) and 16S rRNA bases (yellow). 23S rRNA is shown in blue. **c**) G530 of 16S rRNA in the doublet-decoding structure with open 30S conformation. G530 is withdrawn from the decoding center. Colors as in b, mRNA-GCA in green. Insets show zoomed-out view of the respective ribosome structure in the same orientation.
